# *Populus* VariantDB v3.2 facilitates CRISPR and Functional Genomics Research

**DOI:** 10.1101/2025.04.20.649720

**Authors:** Ran Zhou, Sakshi R. Seth, Jacob Reeves, Andrew H. Burns, Chen Hsieh, Thomas W. Horn, Liang-Jiao Xue, Chung-Jui Tsai

**Affiliations:** Warnell School of Forestry and Natural Resources, University of Georgia, Athens, GA 30602, USA; Department of Genetics, University of Georgia, Athens, GA 30602, USA; Institute of Bioinformatics, University of Georgia, Athens, GA 30602, USA; School of Computing, University of Georgia, Athens, GA 30602, USA; State Key Laboratory of Tree Genetics and Breeding, College of Forestry, Nanjing Forestry University, Nanjing, Jiangsu, 210037, China; Department of Plant Biology, University of Georgia, Athens, GA 30602, USA

**Keywords:** gene editing, sequence polymorphisms, guide RNA, primer, heterozygosity

## Abstract

The success of CRISPR genome editing studies depends critically on the precision of guide RNA (gRNA) design. Sequence polymorphisms in outcrossing tree species pose design hazards that can render CRISPR genome editing ineffective. Despite recent advances in tree genome sequencing with haplotype resolution, sequence polymorphism information remains largely inaccessible to various functional genomics research efforts. The *Populus* VariantDB v3.2 addresses these challenges by providing a user-friendly search engine to query sequence polymorphisms of heterozygous genomes. The database accepts short sequences, such as gRNAs and primers, as input for searching against multiple poplar genomes, including hybrids, with customizable parameters. We provide examples to showcase the utilities of VariantDB in improving the precision of gRNA or primer design. The platform-agnostic nature of the probe search design makes *Populus* VariantDB v3.2 a versatile tool for the rapidly evolving CRISPR field and other sequence-sensitive functional genomics applications. The database schema is expandable and can accommodate additional tree genomes to broaden its user base.

## Introduction

The paradigm-shifting CRISPR technology has enabled targeted genome editing with unprecedented efficiency in many non-model species, including woody perennials (Tsai and Xue 2015; Bewg et al. 2018; Goralogia et al. 2021; Huang et al. 2022; Anders et al. 2023). This powerful technology relies on guide RNAs (gRNAs) that direct CRISPR-associated (Cas) proteins to specific target sites for cleavage, binding, or other effector-assisted activities. Consequently, the precision of CRISPR on-target activities and the minimization of off-target effects depend directly on gRNA design. Various tools have been developed to facilitate gRNA design in plant genome editing (Lei et al. 2014; Xie et al. 2014; Stemmer et al. 2015). Initially limited to model species, some of these gRNA design programs have now expanded to cover tree genomes such as *Citrus, Eucalyptus, Malus*, and *Populus* (Stemmer et al. 2015; Liu et al. 2017). Some programs also support gRNA design for new Cas proteins or variants with different PAM (protospacer adjacent motif) recognition sequences.

Major limitations to existing gRNA design tools include outdated genome versions and the use of polymorphism-blind consensus genomes as references. Sequence polymorphisms are common in outcrossing species, hybrids, or polyploids, and they can affect precision and efficiency of CRISPR experiments (Zhou et al. 2015). This limitation arises from genome assembly practices that typically produce a consensus genome per diploid species as the end product (Sedlazeck et al. 2018). Recent advances in long-read sequencing technologies have now made haplotype-resolved genome assemblies routine (Zhang et al. 2021; Zhou et al. 2023; Carey et al. 2024). The inclusion of haplotype genomes in gRNA design can improve accuracy but complicate the design workflow due to the need to crosscheck multiple genomes. Beyond gRNA design, primers for downstream mutation pattern determination by PCR or amplicon deep-sequencing are also sensitive to sequence polymorphisms. However, most primer design software accepts only a single sequence as input, necessitating additional inspections to ensure their specificity or multiplicity.

The *Populus* VariantDB database was initially created to aid gRNA and primer design for the transformation model *P. tremula* × *alba* INRA 717-1B4 (hereafter 717) before its genome was sequenced (Zhou et al. 2015). We used 717 resequencing data to call variants against the *P. trichocarpa* reference and generated a variant-substituted 717 (s717) custom genome (Xue et al. 2015) to build the first version of the VariantDB. While this approach was effective for variant-aware gRNA design, noncoding sequence divergence and copy number (including presence-absence) variation between genotypes could not be resolved, hindering mutation pattern determination (Bewg et al. 2022; Chen et al. 2023). Here, we present Variant DB v3.2 based on the recently released 717 genome with chromosome-scale assembly of its two haplotypes. Improvements include annotated genomic features, such as coding or putative promoter regions, and improved search speed. We also integrated RazerS3 (Weese et al. 2012) to allow gaps in mapping to improve analysis sensitivity of noncoding regions like promoters and introns where insertion-deletion (indel) polymorphisms are frequent. Finally, the database schema is expandable and can accommodate additional genomes to broaden its user base. The platform-agonistic nature of the probe search design makes *Populus* VariantDB v3.2 a versatile tool to support CRISPR and functional genomics applications in multiple poplar species.

## Materials and Methods

### System architecture and design

JavaScript was used for the development of an interactive and dynamic web experience. The user interface front-end of VariantDB was developed with the open-source JavaScript framework Vue.js. The back-end application was integrated with multiple bioinformatic tools for efficient data handling and read mapping analysis. VariantDB is currently hosted on an x86_64 architecture machine with Intel(R) Xeon(R) Gold 6130 CPU at 2.10 GHz (two cores) running Ubuntu 22.04.5 LTS. The VariantDB search is executed by calling a JavaScript file from the command line. The script reads the user input and checks if it is a valid DNA sequence to be searched against the specified genome databases. The sequence search is implemented using either BatMis v3 (Tennakoon et al. 2012) when the input length is less than 50 mers, Bowtie2 (Langmead and Salzberg 2012) when the input is longer than 50 mers, or RazerS3 (Weese et al. 2012) when gapped alignment is requested. BatMis (Basic Alignment tool for Mismatches) is a Burrows– Wheeler Transform-based aligner for fast short-read mapping while effectively handling multiple mismatches (Tennakoon et al. 2012). RazerS3 is a sensitive short-read aligner that scores mismatches and indels equally as errors in edit distance, unlike Hamming distance used in other aligners that only allows replacements and ignores indels (Weese et al. 2012). Once the input sequence is validated (only standard bases, A, T, C, and G, are accepted), a search is processed with the specified aligner, genome(s), and parameters as follows: 1) Batmis: with -m50, 2) Bowtie2: with -k 30 --very-sensitive --no-hd, and 3) RazerS3: with -i for identity percentage calculated from the user-specified mismatch number. Once the search process is completed, the output of hits is parsed into a list of intervals. The overlaps between hit intervals and any annotated features of the genome are extracted using BEDTools (Quinlan and Hall 2010). Select features, such as CDS and promoter, are highlighted in color as part of the output rendering process, which is coded in the main application for web display.

### Preparation of genome index and annotation files

The genome sequences and associated annotation files of *P. trichocarpa* Niqually-1, *P. deltoides* WV94, and *P. tremula* × *alba* INRA 717-1B4 (717) were downloaded from Phytozome (https://phytozome-next.jgi.doe.gov/) (Goodstein et al. 2012) or provided by Shawn Mansfield (University of British Columbia, Canada) in the case of *P. alba* × *grandidentata* P39. For web display, the chromosome identifiers were simplified to Chr01 to Chr19, but the names of unanchored scaffolds were kept as is. For 717 and P39, the two haplotype genomes were merged into one and their chromosome identifiers were renamed to A01-A19, G01-G19, and T01-T19 for the *P. alba, P. grandidentata*, and *P. tremula* subgenome, respectively. The BatMis genome index files were created by running “build_index genome_file_name” according to the user manual (https://code.google.com/archive/p/batmis/wikis/User_Manual.wiki) (Tennakoon et al. 2012). No genome indices are required for RazerS3.

The standard gff or gff3 annotation files were used to retrieve genomic features containing chromosome (or scaffold) names, feature types (CDS, intron, 5’-UTR, and 3’-UTR), start positions, and strand orientation. The complement function of BEDTools (Quinlan and Hall 2010) was used to process intergenic intervals. We then designated featureless hits within 1 kb upstream of an annotated gene on the positive strand (or downstream if on the negative strand) as potential promoter regions.

## Results and Discussion

### User-friendly capability for gRNA or primer query

The *Populus* VariantDB was developed to enable easy assessment of sequence polymorphisms in the target regions of gRNAs and primers designed for 717, a hybrid poplar commonly used in transgenic research. To maximize flexibility and adaptability in the rapidly evolving CRISPR field, we intentionally avoided duplicating efforts with existing software programs or algorithms for gRNA and primer design. Researchers can use any program of their choice for the design and then submit the resulting short sequences to query VariantDB with one or more genomes (Figure 1A).

**Figure 1.**
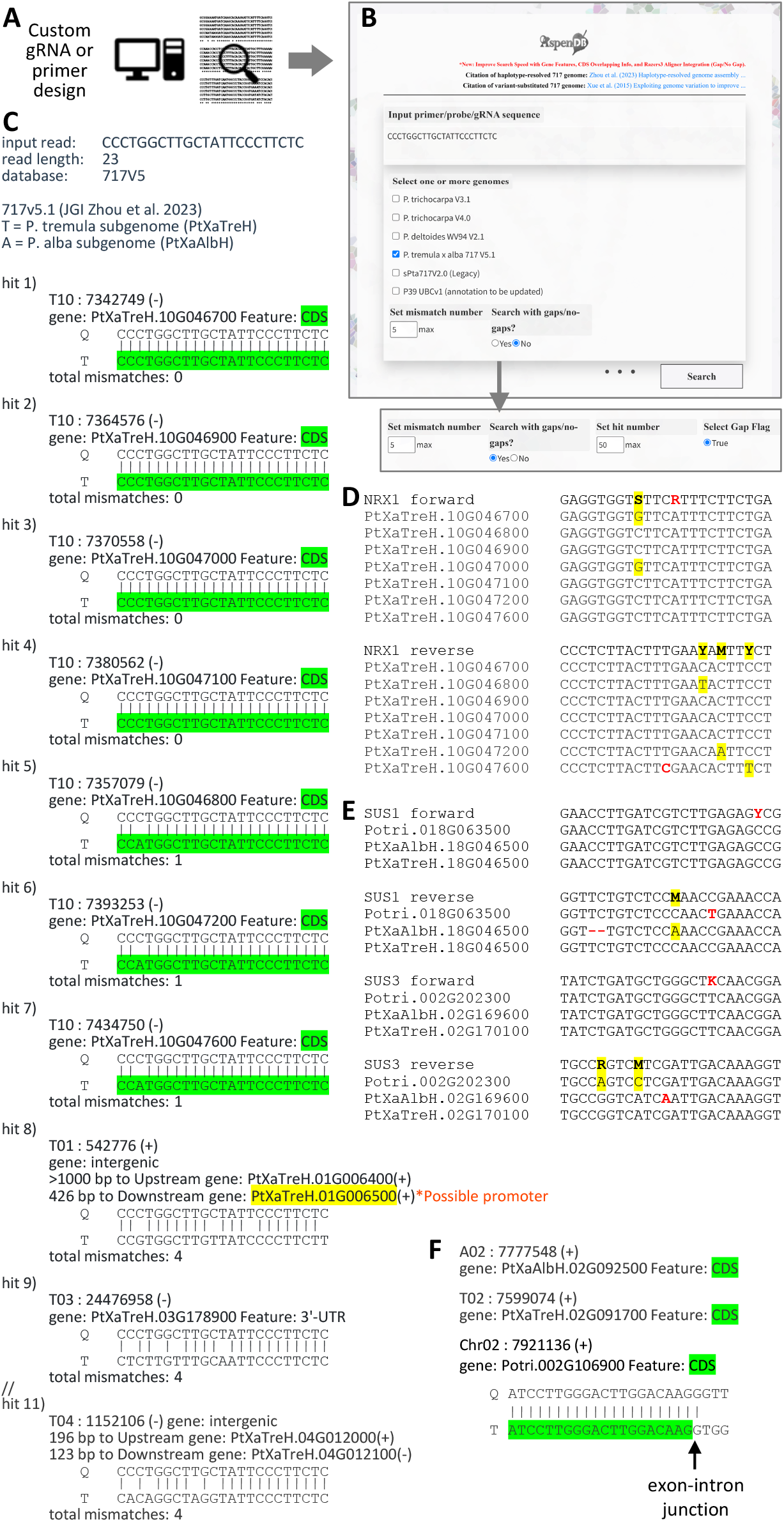
Schematics of VariantDB and representative applications. (A)Users-supplied gRNA or primer sequences can be queried against the database. (B) The database front-end with customizable parameters, either allowing no gaps (top) or with gaps (bottom). (C) A representative output using *NRX1* gRNA as query against the JGI 717 v5.1 genome. CDS and putative promoter are color-coded. (D) Alignments of previously designed *NRX1* amplicon sequencing primers with VariantDB outputs from 717 v5.1. Wobble bases and SNPs are shaded in yellow and mismatches are shown in red. (E) Alignments of previously designed qRT-PCR primers with VaraintDB outputs from *P. trichocarpa* v4 and 717 v5.1. The *P. alba* allele with 2-nt deletions at the *SUS1* reverse primer target site was obtained by allowing gaps in the search. (F) Consolidated hits from *P. trichocarpa* v4 and 717 v5.1 genomes for a previous primer spanning exons. Arrow indicates the exon-intron junction at the target site. The full list of primers is provided in Table S1.

Customizable parameters include the maximum number of mismatches (default: 5) and the option to allow gaps (default: no) (Figure 1B). When the option for gapped alignments is selected, the gap flag is triggered, allowing users to set the mismatch number (default: 5) and hit number (default: 50) (Figure 1B). The output provides a ranked list of genome hits, from best (perfect match) to worst (highest number of mismatches). Each hit is identified with the mapped chromosome, start position, and strand orientation, with an A or T prefix on the chromosome number for *alba* or *tremula* subgenome, respectively (Figure 1C). The corresponding gene models and annotated features, such as CDS, intron, 5’-UTR, 3’-UTR, or putative promoter, are also provided, with some highlighted in color (Figure 1C). For intergenic hits, the upstream and downstream gene models, distances, and strand orientations are shown. Alignments between the query (Q) and target (T) sequences are displayed. The results are not saved in the database, but users can copy and paste the output in text format for their records or modify it for display as shown in Figure 1C.

### Case studies of genes with copy number variation between genotypes

Copy number variations between the 717 and *P. trichocarpa* genomes posed challenges in previous CRISPR studies using the variant-substituted custom genome s717 v2 (Bewg et al. 2022; Chen et al. 2023). For example, the small redox protein nucleoredoxin 1 (NRX1) is encoded by a tandem array of eight genes on Chr10 of *P. trichocarpa* (Chen et al. 2023) but the copy number varied from seven in *P. tremula* to zero in *P. alba* based on the JGI v5.1 717 genome (Zhou et al. 2023). A consensus gRNA designed based on s717 v2 was later confirmed with the JGI v5.1 717 genome to indeed target variant-free regions of the NRX1 tandem duplicates (Chen et al. 2023). However, initial amplicon sequencing primers designed based on s717 v2 were not ideal. The forward primer included an unnecessary wobble at an invariable position, and the reverse primer missed an SNP (Figure 1D). This resulted in no amplification of *PtaNRX1*.*7*, an issue that remained unknown until the 717 v5.1 genome became available. This example serves to illustrate how mismatches in gRNA or primer sequences can affect experimental outcomes, and troubleshooting these issues can be difficult and time-consuming.

The VariantDB is also valuable for vetting published RT-qPCR primers, which often span the 3’-UTR for maximal discrimination between highly homologous genes. Like gRNAs, the specificity of primers is influenced by sequence and copy number polymorphisms of the study organism. This is especially true for primers designed based on experimentally cloned cDNAs, ESTs, or the *P. trichocarpa* reference genome with limited or no haplotype information. We tested 39 primers for 25 genes involved in sugar metabolism from a previous study (Payyavula et al. 2011). These primers were designed based on the older *P. trichocarpa* genome v2 and cross-checked against *Populus* ESTs to include wobbles for polymorphic bases. Despite this practice, only 16 primers match perfectly with the target genes in the *P. trichocarpa* v4 and 717 v5.1 genomes and 8 additional primers were predicted to be functional but harbored unnecessary wobbles (Figure 1E, Supplemental Table S1). The remaining 15 primers had 1-3 mismatches, including indels (Figure 1E, Supplemental Table S1). As a result, these primers were predicted to work effectively for only 28 of the 50 *P. tremula* and *P. alba* alleles in 717 versus 21 of the 25 *P. trichocarpa* genes. We acknowledge that primers with mismatches have been shown to work, albeit with reduced efficiencies (Gibbs et al. 1989; Simsek and Adnan 2000). Regardless, these findings suggest that primers designed without haplotype information should be used with caution and that VariantDB can facilitate primer validation. It should be noted that primers spanning exons are not displayed correctly against the current genomic database (Figure 16F) and will require manual curation.

### Expansion and limitations

The database schema is readily expandable to accommodate additional genomes. Currently, the *Populus* VariantDB hosts the genomes of *P. trichocarpa* Nisqually-1 v3.1 and v4.0 (Tuskan et al. 2006) and *P. deltoides* WV94 v2.1 from Phytozome v13 (Goodstein et al. 2012), as well as the haplotype-resolved *P. alba* × *grandidentata* P39 v1 (provided by Shawn Mansfield, personal communication), in addition to the 717 JGI v5.1 (Zhou et al. 2023). The first two are consensus genomes with no polymorphism information but can be easily upgraded when haplotype-resolved versions become available in the future.

Known limitations include erroneous annotation of genomic features. The annotation quality of a genome is known to evolve with advancements in sequencing technologies, genome assembly methods, computational gene calling algorithms, and experimentally validated genes. Regular updates of the genomes will ensure that the database incorporates the most current genome assembly and annotation versions. Another limitation is the inability to handle exon-spanning sequences, such as RT-qPCR primers, as discussed above (Figure 1F). This could be alleviated by including transcriptomes as additional references. Finally, while the search speed has improved in the current implementation, with most single-genome searches completed within seconds, searches against multiple genomes or with the option of allowing gaps are slower due to the available computing resources. We suggest using the no-gap option for most applications, and only testing the more computationally demanding option of allowing gaps when the search does not return the expected results. For instance, the missing *P. tremula* or *P. alba* allele in the standard 717 search output was recovered by allowing gaps, as illustrated for the *SUS1* (sucrose synthase) and *SPS6* (sucrose phosphate synthase) reverse primers due to indels (Figure 1E, Supplemental Table S1).

In conclusion, the updated VariantDB v3.2 offers a unique genomic resource that supports *Populus* functional genomics research. It complements existing genomic databases and CRISPR gRNA and primer design programs by providing easily accessible sequence polymorphisms of heterozygous genomes that can affect experimental outcomes. The database can be expanded to include additional tree genomes to benefit more researchers.

## Acknowledgements

The authors thank current and former lab members for their valuable input to the development of VariantDB, the Joint Genome Institute and the HudsonAlpha Institute for Biotechnology teams for *P. trichocarpa* Nisqually-1, *P. deltoides* WV94, and *P. tremula* × *alba* genomes, and Shwan Mansfield and Qian Wang for the *P. alba* × *grandidentata* P39 genome.

## Authors’ contributions

C.J.T. conceived the idea, R.Z., S.R.S., J.R., L.-J.X., A.H.B., C.H., and T.W.H. built and improved the database, R.Z. drafted the manuscript, C.-J.T. revised the manuscript.

## Supplementary data

Table S1. List of published qRT-PCR primers and their VariantDB matches

## Data availability statement

VariantDB source code is available on github (https://github.com/TsailabBioinformatics/ProbeSearchV3).

